# Myomegalin regulates Hedgehog pathway by controlling PDE4D at the centrosome

**DOI:** 10.1101/2020.04.24.059923

**Authors:** Hualing Peng, Jingyi Zhang, Amanda Ya, Winston Ma, Sammy Villa, Shahar Sukenik, Xuecai Ge

## Abstract

Mutations in the Hedgehog (Hh) signaling are implicated in birth defects and cancers, including medulloblastoma, one of the most malignant pediatric brain tumors. Current Hh inhibitors face the challenge of drug resistance and tumor relapse, urging new insights in the Hh pathway regulation. Our previous study revealed how PDE4D controls global levels of cAMP in the cytoplasm to positively regulate Hh signaling; in the present study we found that a specific isoform PDE4D3 is tethered to the centrosome by myomegalin, a centrosome/Golgi associated protein. Myomegalin loss dislocates PDE4D3 from the centrosome, leading to local PKA over-activation and inhibition of the Hh signaling, leaving other PKA-related pathways unaffected. Myomegalin loss suppresses the proliferation of granule neuron precursors, and blocks the growth of medulloblastoma in mouse model. Our findings specify a new regulatory mechanism of the Hh pathway, and highlight an exciting therapeutic avenue for Hh-related cancers with reduced side effects.

## Introduction

The Hedgehog (Hh) pathway is widely implicated in birth defects and human tumors(Briscoe and Therond, 2013). One of the Hh-related tumors is medulloblastoma, a malignant pediatric brain tumor(Goodrich *et al.*, 1997). Current treatment of medulloblastoma, surgery removal followed by chemo- or radiotherapy, brings devastating side effects to the young patients(Fouladi *et al.*, 2005); while the available Hh-pathway inhibitor targeting Smoothened (Smo) is challenged by drug resistance and tumor relapse(Yauch *et al.*, 2009). Therefore, new approaches to inhibit Hh signaling are needed. The Hh signal transduction involves a series of protein transport into and out of the primary cilium, and eventually converges on the regulation of Gli transcription factors(Hui and Angers, 2011). Without the ligand Sonic Hedgehog (Shh), the receptor Patched (Ptch) resides in the cilium and prevents the cilium translocation and activation of Smo. Upon Shh stimulation, Ptch exits the cilium, followed by Smo’s accumulation and activation in the cilium. The signaling cascade ultimately activates the transcription activator Gli2, and eliminates the transcription suppressor Gli3R, a proteolytic product from the Gli3 full length (FL) protein(Wang and Li, 2006; Han and Alvarez-Buylla, 2010). The activated Hh signaling quickly induces the transcription of Gli1, an amplifier of Hh signaling, forming a positive feedback loop.

PKA plays a central role in Hh signaling activation and Gli regulations. PKA phosphorylates Gli3, which primes its further phosphorylation by GSK and CK1. The phosphorylated Gli3 was recognized by the ubiquitin proteosome system that cleaves Gli3FL into Gli3R(Wang and Li, 2006). In addition, PKA also controls Gli2 activation. Within the cell, PKA concentrates at the centrosome (cilium base) where it controls the cilium translocation of Gli2, a step required for Gli2 activation(Tuson, He and Anderson, 2011). Genetic removal of PKA leads to full activation of Hh pathway in the developing neural tube(Epstein *et al.*, 1996; Huang, Roelink and McKnight, 2002; Tuson, He and Anderson, 2011), further substantiating the strong inhibitory effect of PKA on Hh signaling. Recent studies from Mukhopadhyay lab suggest that inhibiting the cAMP-PKA levels in the cilium markedly activates Hh signaling in a manner independent of Smo activation (Somatilaka *et al.*, 2020). Conversely, pharmacological activation of PKA inhibits Hh signaling and suppresses Hh-related tumor growth(Yamanaka *et al.*, 2010, 2011). However, PKA is widely involved in many signaling and metabolic pathways; ubiquitous activation of PKA inevitably impacts all signaling pathways. Hence, treatments directly targeting PKA are not practical due to their severe side effects. To avoid these side effects, one feasible strategy is to selectively control PKA activities at the specific subcellular sites where it regulates Hh signaling, leaving other pathways unaffected.

It is known that PKA activity in the cell is compartmentalized by forming complexes that include cAMP-specific phosphodiesterase (PDE) (Zaccolo and Pozzan, 2002; Houslay, 2010; McCormick and Baillie, 2014). In specific compartments, PKA activity is precisely regulated by PDE. In our previous studies, we found that PDE4D, recruited to the cytoplasmic membrane by sema3-Neuropilin signaling, governs cAMP levels in the entire cell to regulate Hh signaling(Tyler Hillman *et al.*, 2011; Ge *et al.*, 2015). Our results were corroborated by Williams et al. who independently discovered PDE4D as a positive regulator of the Hh pathway in a chemical screen(Williams *et al.*, 2015). Since PKA at the centrosome directly participate in Hh signaling(Barzi *et al.*, 2010; Tuson, He and Anderson, 2011), can we selectively manipulate PDE4D activity at the centrosome to control local PKA activity? In the current study, we found an approach to dislocate PDE4D3 from the centrosome; the subsequent elevation in local PKA activity suppresses Hh signal transduction and Hh-related tumor growth. Our results highlight an exciting avenue to treat Hh-related cancers with reduced side effects.

## Results

### Myomegalin (Mmg) interacts with PDE4D3 at the centrosome

To identify an effective approach of selectively modulating cAMP levels at the centrosome, we did a literature search on the subcellular localization of all cAMP-specific phosphodiesterases. We found that one PDE4D isoform, PDE4D3 was reported to interact with Mmg, a protein associated with the centrosome/Golgi(Verde *et al.*, 2001). But it remains unclear whether PDE4D3 localizes to the centrosome and whether it is involved in the regulation of the Hh signaling. To answer these questions, we first validated the Mmg-PDE4D3 interaction. Mmg is a large protein of 270KD, and the full-length protein is not effectively expressed in cells. But in the previous study the C-terminus of Mmg was identified to mediate its interaction with PDE4D3(Verde *et al.*, 2001) (Fig. 1A). We thus fused this domain (Mmg-C) with Flag and expressed it together with HA-PDE4D3 in the cell. We then performed co-immunoprecipitation assay with Flag- and HA- conjugated magnetic beads, and found that two proteins co-immunoprecipitated each other (Fig. 1B). It is noteworthy that although Flag antibody pulled down significant amount of Flag-Mmg (red triangle in Fig. 1B), it only coimmunoprecipitated small amount of HA-PDE4D3 (red star in Fig. 1B), presumably because only a small fraction of HA-PDE4D3 in the cell is interacting with Mmg. This is consistent with what we observed in immunostaining results in Fig. 1E.

**Figure 1.**
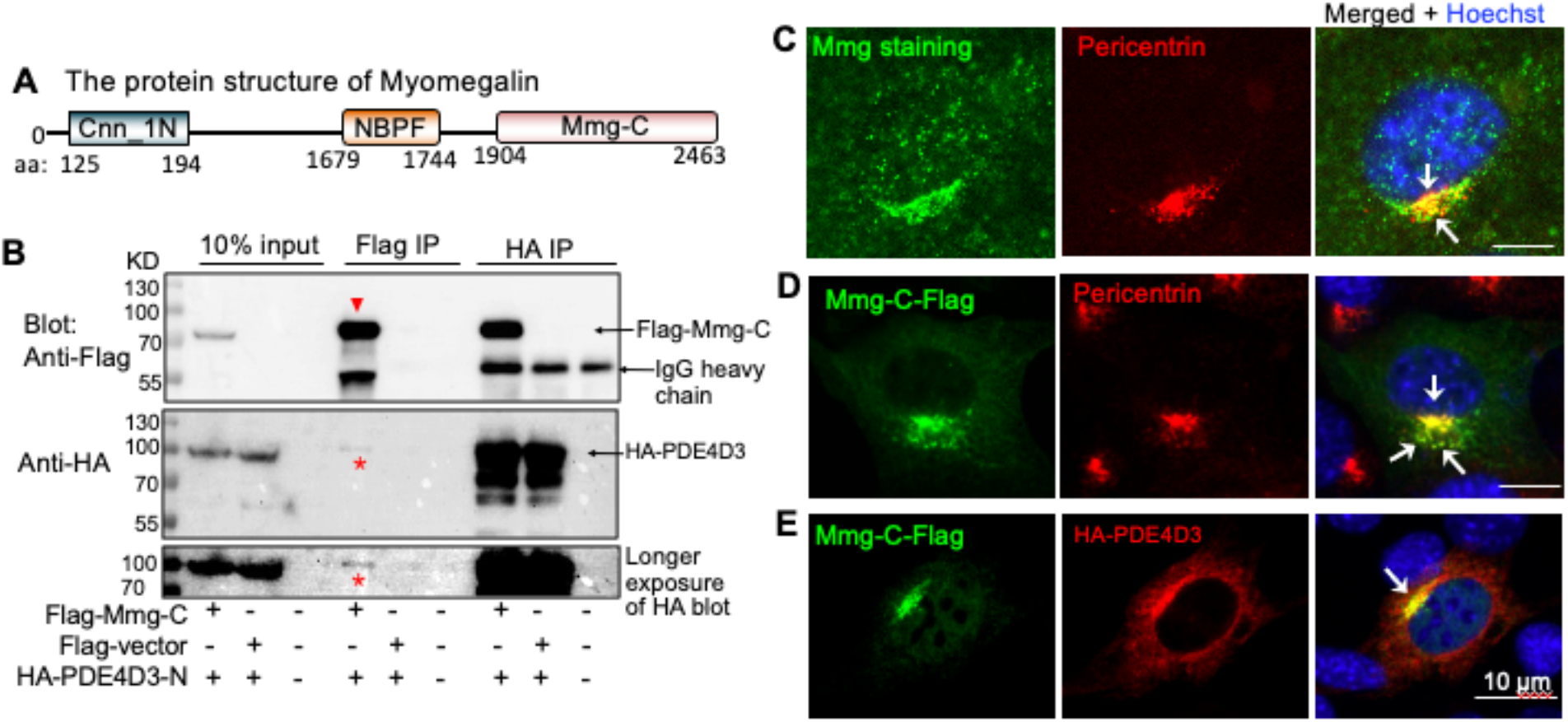
PDE4D3 interacts with Mmg at the centrosome. (A) Protein structure of Mmg. Cnn_1N, Centrosomin N-terminal motif 1; NBPF (DUF1220), domain of Neuroblastoma breakpoint family; Mmg-C, the domain previously shown to interact with PDE4D3. (B) When overexpressed in HEK293T cells, HA-PDE4D3 and Flag-Mmg C-terminus co-immunoprecipitated with each other, suggesting the interaction of the two proteins. (C) Immunostaining of endogenous Mmg shows that Mmg overlaps with pericentrin, a marker for the centrosome and pericentriolar materials (white arrows). (D) When expressed in NIH3T3 cells, Mmg-C colocalizes to the centrosome and pericentriolar material (white arrows). (E) When expressed in NIH3T3 cells, HA-PDE4D3 diffusively distribute to the cytoplasm, but a significant fraction of PDE4D3 is recruited by Mmg to the centrosome and pericentriolar material (white arrows).

To validate the subcellular localization of Mmg, we stained NIH3T3 cells with the Mmg antibody. As reported before(Roubin *et al.*, 2013), Mmg immunofluorescence significantly overlaps with pericentrin, a marker of the centrioles and pericentriolar material (Fig. 1C). The non-centrosomal Mmg signal may represent its localization to Golgi. Mmg-C exhibits similar localization pattern when expressed in NIH3T3 cells (Fig. 1D). We then expressed both HA-PDE4D3 and Flag-Mmg-C in the cell. HA-PDE4D3 overlaps with Flag-Mmg at the centrosome/Golgi area, although a significant fraction of HA-PDE4D3 also diffusively distributes to the cytosol (Fig. 1E). Taken together, these results suggest that Mmg may recruit a small fraction of PDE4D3 from the cytosol to the centrosome.

### Mmg loss impairs Hh signal transduction and dislocates PDE4D3 from the centrosome

Next, we tested whether eliminating PDE4D3 from the centrosome impacts the Hh pathway. We silenced Mmg expression with shRNA in NIH3T3 cells, a cell line that contains all components of the Hh pathways and is commonly used to study Hh signaling transduction. Two of the five tested shRNAs significantly reduced the transcript and the protein levels of Mmg (Fig. S1A-B). We then treated cells with SAG, a small molecule agonist of the Hh pathway, and assessed the Hh pathway activation with qPCR measuring the transcript level of the Hh target gene Gli1. Mmg shRNA significantly reduced SAG-induced Gli1 expression, indicating that Hh signal transduction was impaired (Fig. S1C).

To thoroughly eliminate Mmg protein expression, we employed CRISPR/Cas9 to knockout Mmg in mouse embryonic fibroblasts (MEFs). We choose MEF because it transduces Hh signaling but has lower ploidy level than NIH3T3 cells. We used two gRNAs targeting the 1^st^ exon of Mmg, and transfected the plasmid containing the two gRNAs and Cas9 into MEF cells, together with EGFP. Single cell clones were isolated via flow cytometry and expanded (Fig. 2A). We obtained two cell clones (#7, #10) of Mmg knockout (KO). Both clones appear normal in cell morphology and cell proliferate (data not shown), and the Mmg mRNA and protein levels are undetectable (Fig. 2B-C). Interestingly, among the 4 alternative splicing isoforms of mouse Mmg (https://www.ncbi.nlm.nih.gov/gene/83679), CRISPR/Cas9 abolished the expression of the longer isoforms (~270KD) and spared the shorter isoform (~130KD) (Fig. 2C), presumably because the shorter isoform uses an alternative transcription starting point. The shorter isoform, however, does not interact with PDE4D3 as it lacks the C-terminus. To identify the INDEL mutations induced by CRISPR/Cas9, We amplified exon 1 and its flanking region with PCR from Mmg KO cells, and sequenced individual PCR products. The sequencing results show that 3 type of mutations were generated in each clone, resulting in frameshift that eventually leads to nonsense-mediated mRNA decay (Fig. S2A-C).

**Figure 2.**
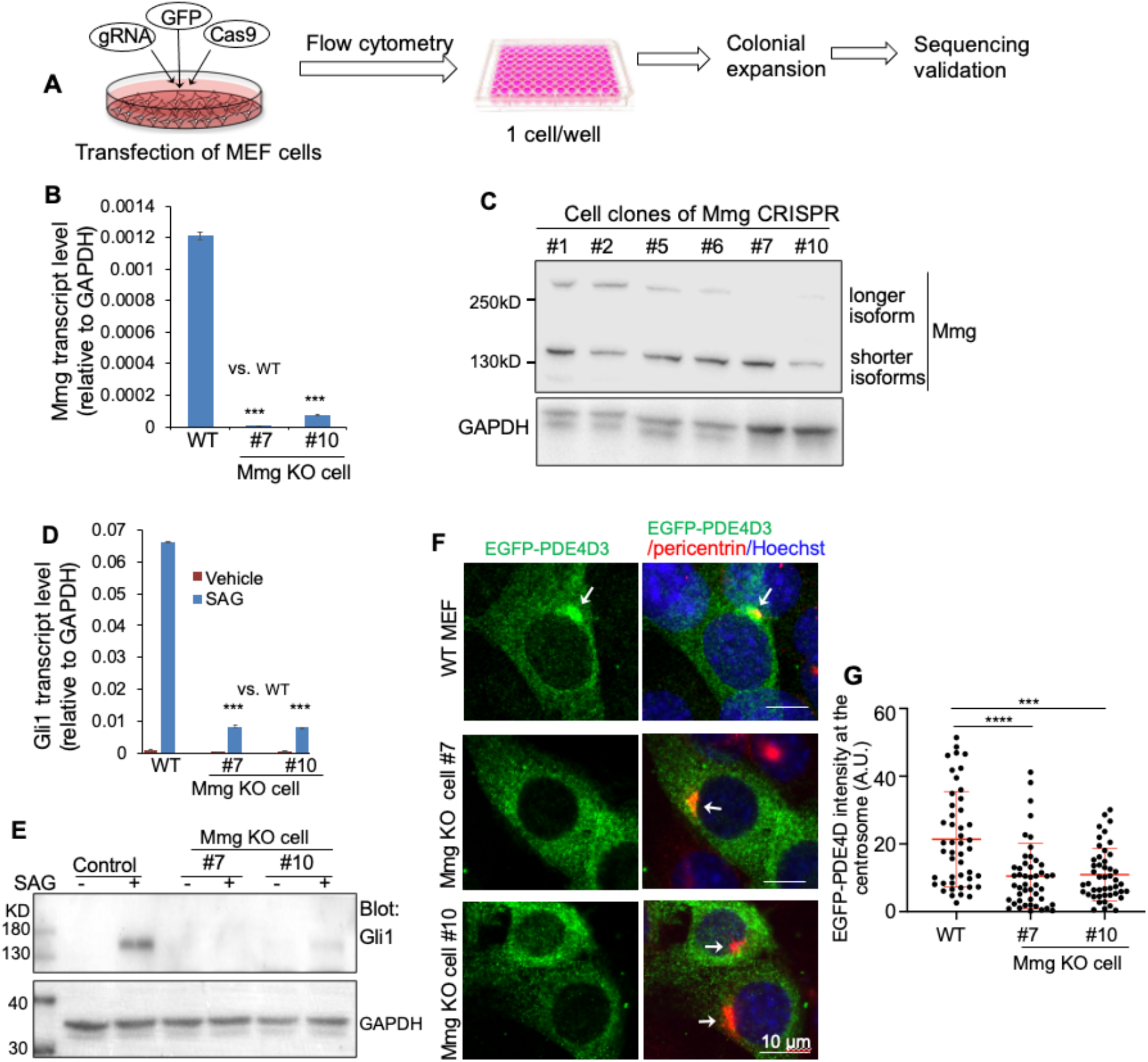
Mmg knockout dislocates PDE4D from the centrosome and impairs Hh signal transduction. (A) Procedure of generating Mmg CRISPR cell clones. 30 cell clones were established and tested for Mmg transcript levels and protein levels. (B) In two of the Mmg CRISPR cell clones, the transcript of Mmg was hardly detectable by qPCR. (C) Western blot shows that in cell clone #7 and #10, the CRISPR abolished the expression of the longer isoforms of Mmg, but the shorter isoform remains unaffected. (D-E) Mmg CRISPR knockout clones were stimulated with SAG for 24hr, and Hh signaling activity was evaluated by Gli1 transcript levels and protein levels. The Hh activity was dramatically suppressed in Mmg knockout cells. (F) Representative images of EGFP-PDE4D3 expressed in wild type or Mmg knockout cell clones. PDE4D3 concentrates at the centrosome and pericentriolar material in wild type cells (white arrow); however, in CRISPR cell clones, it only exhibits diffusive distribution to the cytoplasm and lacks the significant overlap with pericentrin (white arrows). (G) Quantification of PDE4D intensity at the centrosome. All error bars represent SD; statistics in B and D: Student’s t-Test. **p<0.01, ***p<0.001. Statistics in G: Kruskal–Wallis non-parametric One-Way ANOVA, followed by Dunn’s multiple comparison. ***p<0.001, ****p<0.0001. A.U.: arbitrary unit.

To determine the Hh signaling in Mmg KO clones, we stimulated cells with SAG, and detected Gli1 expression with qPCR and western blot. SAG-induced Gli1 expression was dramatically reduced at the transcript and protein levels in both Mmg KO cell clones (Fig. 2D-E). These results suggest a blockage of Hh transduction after Mmg loss.

Next, we determined the impact of Mmg loss on PDE4D3 localization at the centrosome. Due to the high similarity between PDE4D isoforms, the antibody specific to PDE4D3 is unavailable. Therefore, we expressed very low levels of EGFP-PDE4D3 in Mmg KO cells to mimic the endogenous protein. As expected, in wild type cells PDE4D3 shows significant overlap with pericentrin, in addition to its diffusive localization to other subcellular sites (Fig. 2F). However, in Mmg KO cells, the intensity of PDE4D3 at the centrosome is significantly reduced (Fig. 2F-G). Therefore, without Mmg, PDE4D3 is dislocated from the centrosome.

In summary, our data suggests that loss of Mmg markedly suppresses Hh signal transduction, and dislocates PDE4D3 from the centrosome.

### Mmg loss selectively increases local PKA activity at the centrosome, and blocked further PKA activation by PDE4D inhibitors

Dislocation of PDE4D3 from the centrosome increases local cAMP levels, which may eventually lead to PKA over-activation. To confirm this, we employed two method to evaluate local PKA activity at the centrosome. First, to measure the basal levels of active PKA, we stained cells with an antibody that recognizes active PKA (phosphoPKA T197). This antibody has been used to evaluate PKA activity in previous studies(Barzi *et al.*, 2010; Tuson, He and Anderson, 2011; Ge *et al.*, 2015). We highlighted the centrosome and pericentriolar area with pericentrin staining, and measured the phosphoPKA levels in this area in ImageJ. As expected, the centrosomal active PKA levels are much higher in Mmg knockout cells compared to that in wild type MEF (Fig. 3A-B). Further, expressing exogenous Mmg in Mmg KO cells restored active PKA levels to normal (Fig. 3B). This change of PKA activity, however, is limited locally to the centrosome, since the overall phosphoPKA levels remain the same in Mmg KO cells (Fig. 3C). In addition, we detected the phosphorylation levels of CREB, the cytosolic substrate of PKA(Shaywitz and Greenberg, 1999). The phospho-CREB (S133) levels show no difference between wild type and Mmg knockout cells (Fig. 3C). Thus, Mmg loss increased the basal PKA activity selectively at the centrosome.

**Figure 3.**
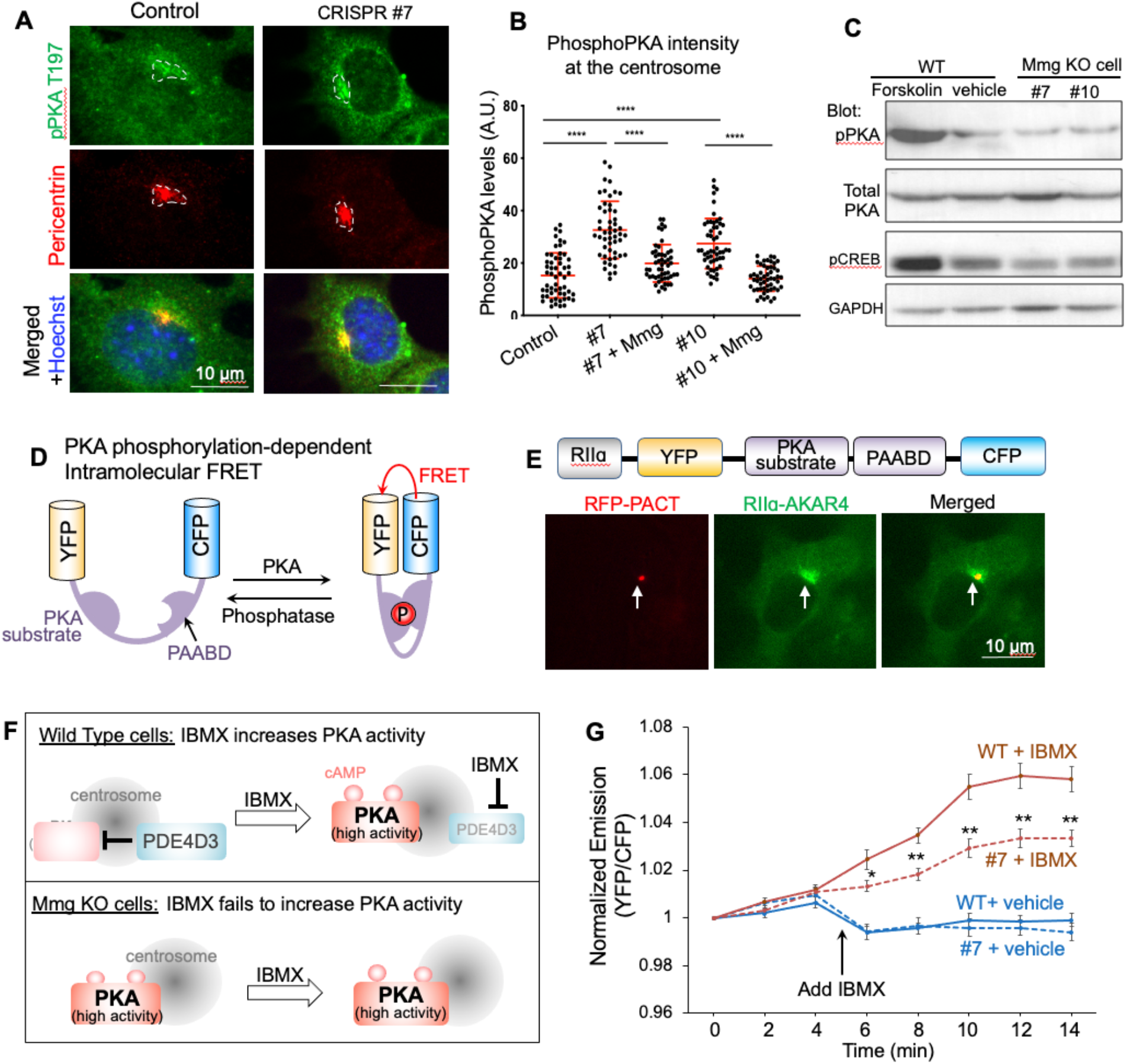
Mmg loss increases basal PKA activity at the centrosome, and abolishes further PKA activation by IBMX. (A) Representative images of active PKA immunostaining demonstrate that the basal active PKA levels at the centrosome increased in Mmg knockout cells (white arrows). Dotted lines circled the areas where pPKA intensity were measured based on the staining of pericentrin. (B) Quantification of basal active PKA levels at the centrosome. Expression of exogenous Mmg restored active PKA levels at the centrosome. Data are shown as mean ± SD. Statistics: Kruskal–Wallis non-parametric One-Way ANOVA, followed by Dunn’s multiple comparison. ***p<0.001, ****p<0.0001. A.U.: arbitrary unit. (C) Western blot shows that the global levels of active PKA do not change in Mmg knockout cells. Forskolin treatment serves as a positive control of PKA overactivation in the entire cell. (D) Schematic view of AKAR4, a FRET based probe for PKA activity. The CFP (Cerulean) and YFP (cpVE172) is linked by a linker sequence that contain a PKA phosphorylation site and a phosphoanimo acid binding domain (PAABD). PKA phosphorylation induces conformational change in the linker, which brings CFP and YFP in close proximity to produce FRET. (E) By fusing AKAR4 to the RII subunit of PKA, we targeted the probe to the centrosome. Immunostaining results confirmed the centrosome localization of RIIɑ-AKAR4. (F) Diagram showing that inhibiting PDE4D increases PKA activity in wild type cells, but has little effect on PKA activity in Mmg knockout cells. (G) Normalized emission of FRET acceptor over donor before and after IBMX 0.1mM. The ratio of YFP/CFP at each time point was normalized to time zero. n = 8-13 cells. Data are shown as mean ± SEM. Statistics: t-Test, between the wild type cells and Mmg knockout cells at the same time point. *p<0.05, **p<0.01.

Second, we monitored the dynamic PKA activity in live cells with A kinase-activity reporter (AKAR4), a fluorescence resonance energy transfer (FRET)-based PKA probe developed in Jin Zhang’s lab(Zhang *et al.*, 2001; Herbst, Allen and Zhang, 2011). In this probe, a FRET pair (CFP and YFP) is connected by a linker sequence that contains PKA phosphorylation sites and a phosphoamino acid binding domain (PAABD). PKA phosphorylation induces conformational changes in the linker, which brings the FRET pair in close proximity to efficiently produce FRET (Fig. 3D). To target AKAR4 to the centrosome, we fused it with the regulatory subunit of PKA (PKARIIɑ), a protein predominantly localized to the centrosome(Zhang *et al.*, 2001). As expected, when RIIɑ-AKAR4 was expressed in MEF, the probe is concentrated at the centrosomal area (Fig. 3E). The centrosome is marked by co-expression of RFP-PACT(Gillingham and Munro, 2000).

The PDE inhibitor IBMX is commonly used to enhance the PKA activity, because it effectively elevates cAMP levels in the cell (Zhang *et al.*, 2001; Herbst, Allen and Zhang, 2011). Since PDE4D3 is dislocated from the centrosome in Mmg KO cells and the local PKA activity is constitutively high, we hypothesize that IBMX’s effect will be masked at the centrosome in Mmg KO cells (Fig. 3F). To test this hypothesis, we treated cells with IBMX and analyzed FRET locally at the centrosome. The FRET efficiency was analyzed as the ratio of YFP/CFP, and this ratio at each time point was normalized to time 0 (Fig. 3G & Fig. S3). In WT cells, IBMX gradually increased FRET efficiency, peaking at 12 min. In contrast, Mmg knockout significantly dampened FRET efficiency at all time points (Fig. 3G). Taken together, Mmg loss dislocates PDE4D3 from the centrosome, thereby promoting the local basal PKA activity that cannot be further elevated by the PDE inhibitor.

### Mmg loss promotes Gli3R production and blocks Gli2 transportation to the cilium tip

The transcription factor Gli2 and Gli3 are PKA substrates in the Hh pathway. After PKA phosphorylation, Gli3 is proteolytically processed into Gli3R, a transcription repressor (Fig. 4A). Upon Hh signaling activation, the Gli3 processing ceases and Gli3R levels markedly reduce (Wang and Li, 2006; Humke *et al.*, 2010; Tukachinsky, Lopez and Salic, 2010; Hui and Angers, 2011). We examined Gli3 processing by western blot. In wild type cells, SAG treatment significantly reduced Gli3R levels in cell lysates. In contrast, in Mmg CRISPR clones, the Gli3R levels remain the same after SAG treatment (Fig. 4B). It is likely that without Mmg, the hyperactive PKA at the centrosome continues to phosphorylate Gli3 to promote Gli3R production even after SAG treatment. PKA affects the proteolysis of Gli2 only very slightly but more dramatically controls its accumulation at cilia tips, a step required for Gli2 activation(Barzi *et al.*, 2010; Tuson, He and Anderson, 2011). We therefore examined the levels of Gli2 at the cilia tips after SAG stimulation. The Gli2 intensity at the cilium tips in Mmg KO cells was significantly lower, compared to that in wild type cells (Fig. 4C). When exogenous Mmg was expressed in Knockout clones, the Gli2 levels at the cilium tips was restored (Fig. 4C-D). Therefore, Mmg loss overactivates PKA at the centrosome, which blocks Gli2 transport and activation in the cilium tip, leading to inhibition of the Hh signaling.

**Figure 4.**
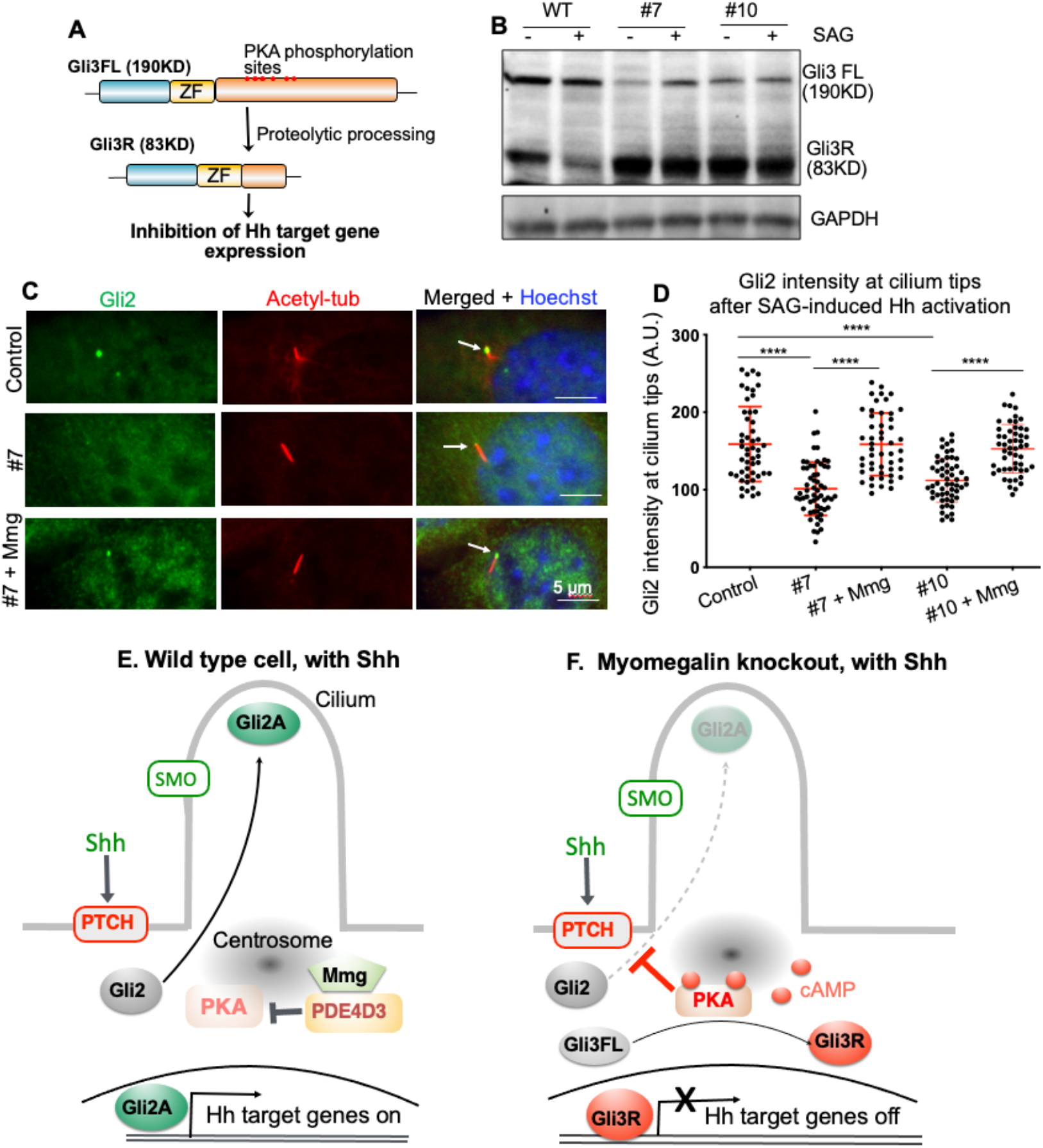
Mmg loss impacts Gli3 processing and Gli2 transportation to the cilium tips. (A) Diagram showing the proteolytic processing of Gli3 after PKA phosphorylation. (B) 24hr SAG treatment reduced Gli3R levels in wild type cells, but not in Mmg knockout cell clones. (C) Representative images of Gli2 immunostaining after cells are stimulated with SAG. White arrows point to Gli2 at the cilium tips. (D) Quantification of Gli2 levels at the cilium tips. Mmg knockout reduced Gli2 levels at the cilium tips, while Mmg overexpression restored Gli2 intensity. Data are shown as mean ± SD. Statistics: Kruskal–Wallis non-parametric One-Way ANOVA, followed by Dunn’s multiple comparison. ****p<0.0001. A.U.: arbitrary unit. (E-F) Diagram showing PED4D3 specifically controls PKA activities at the centrosome to regulate the Hh signaling transduction. Under normal conditions, Upon Shh stimulation, SMO is translocated and activated in the cilium, which then triggers a signaling cascade that reduces cAMP levels at the cilium base. The subsequent inhibition of PKA allows Gli2 to be translocated and activated in the cilium tips (E). Without myomegalin, PDE4D3 is dislocated from the centrosome and fails to degrade the local cAMP. Thus, PKA levels remain high at the centrosome even after Shh stimulations. Hyperactive PKA suppresses Gli2 activation and promotes Gli3R production. As a result, the Hh pathway cannot be activated (F).

In summary, our data suggest a model of how Mmg and PDE4D3 at the centrosome control local PKA activity to regulate the Hh pathway. Without Shh, PKA activity at the centrosome is high due to high local cAMP levels. The centrosomal cAMP may be produced in the cilium by proteins such as GPR16(Mukhopadhyay *et al.*, 2013), or diffuses to the centrosome from other cytosolic areas. After Shh stimulation, GPR161 exits the cilium and stops the cAMP production; the cAMP from nearby cytosolic areas is degraded by PDE4D3. The inactive PKA at the centrosome allows Gli2 to be translocated and activated in the cilium tips, and stops Gli3R production, leading to Hh pathway activation (Fig. 4E). In Mmg KO cells, since PDE4D3 is dislocated from the centrosome, the cAMP diffused from the nearby areas is not effectively degraded and the PKA activity remains high. This suppresses Gli2 activation and keeps Gli3R levels high, and subsequently blocks the activation of Hh signaling (Fig. 4F).

### Mmg loss blocks cell proliferation in primary cultured granule neuron precursors (GNPs)

In the developing cerebellum, Shh is the mitogen that stimulates GNP proliferation(Dahmane and Ruiz i Altaba, 1999; Wallace, 1999; Wechsler-Reya and Scott, 1999). Overactive Hh signaling leads to GNP over proliferation that eventually results as medulloblastoma (MB), one of the most malignant pediatric brain tumor(Kool *et al.*, 2012). In situ hybridization results show that both PDE4D3 and Mmg (www.informatics.jax.org/image/MGI:5332354, www.informatics.jax.org/image/MGI:5333985) are highly expressed in the developing cerebellum(Richter, Jin and Conti, 2005); it is likely that the mechanism of Mmg-PDE4D3 regulation on Hh pathway applies to the control of GNP proliferation. To test this hypothesis, we cultured GNPs from P7 mouse neonates in dishes, and infected GNPs with lentiviral particles expressing shRNA against Mmg (shRNA #99) (Fig. 5A). GNP proliferation was induced by SAG. After 3 days of primary culture, GNP proliferation was assessed by BrdU incorporation assay. As expected, Mmg shRNA significantly reduced Mmg transcript levels, and reduced the rate of BrdU incorporation after pulse labeling (Fig. 5B-D). Hh signal activity was significantly reduced in Mmg knockdown cells, demonstrated by decreased transcript levels of Gli1, a Hh target gene (Fig. 5E). In summary, our results suggest that the same mechanism of Hh signaling regulation by Mmg-PDE4D3 may control GNP proliferation in the developing cerebellum.

**Figure 5.**
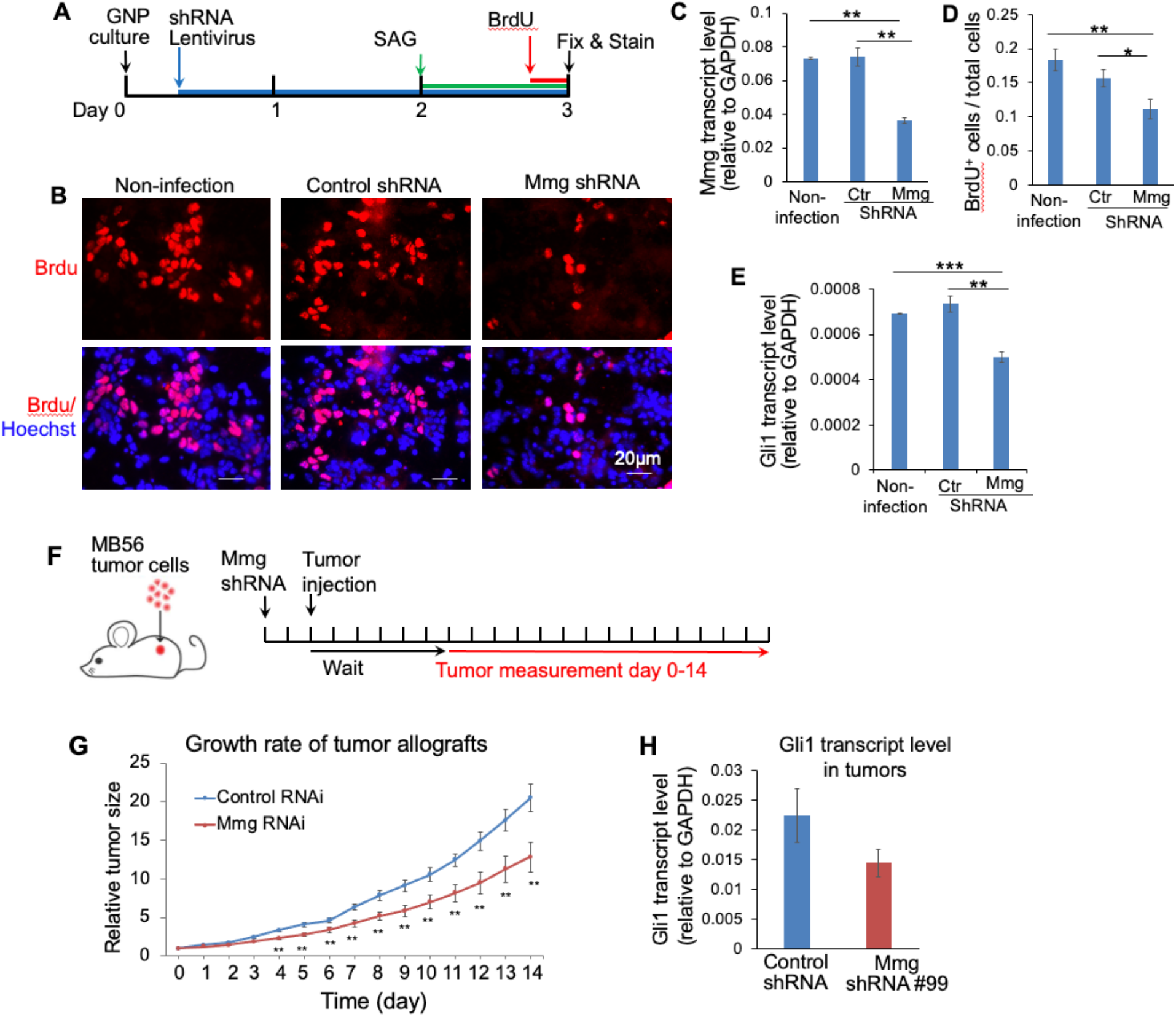
Mmg knockdown blocked cell proliferation in primary cultured GNPs and suppressed the growth rate of medulloblastoma in mouse model. (A) Schematic of BrdU incorporation assay in primary cultured GNPs. Lentivirus expressing shRNA against Mmg or control shRNA were added to the cell 5-6 hr after GNPs are plated in dishes. SAG was added to the culture 24 hr before cells were fixed. BrdU pulse labeling lasted for 4 hr right before cells were fixed. (B) Representative images of BrdU immunostaining in GNPs. (C) Mmg transcript levels at the end of the experiments, measured by qPCR. (D) BrdU incorporation rate in GNPs. (E) Levels of Hh signaling activity evaluated by Gli1 transcript levels. (F) Schematic diagram of the MB56 tumor allograft experiment in mouse. Measurement started 6 day after tumor allograft when the size of tumors could be accurately measured. (G) The relative tumor size is defined as the tumor volume on the indicated day divided by that on day 0. For each treatment 8–9 mice were used, and each mouse was transplanted with two tumors on their hind flank. Results shown are from one of the two independent experiments. (H) At the end of the experiment, the Gli1 transcript levels in 6 of randomly sampled tumors were assessed by qPCR. Hh signaling activity was reduced by Mmg RNAi. Data are presented as Mean ± SEM. Statistics: Student’s t-Test. *p<0.05, **p < 0.01, ***p < 0.001.

### Mmg loss reduced the growth rate of medulloblastoma in mouse model

Next, we assess the effect of Mmg loss on the Hh-related tumor growth in the mouse model of MB subcutaneous xenograft used in our previous studies(Ge *et al.*, 2015). We employed MB56, MB tumor cells directly taken from Ptch^+/−^ mouse, the first and well-established MB mouse model(Goodrich *et al.*, 1997; Purzner *et al.*, 2018). We infected MB56 with lentiviral particles that express Mmg (shRNA #99) or control shRNA. 2 days after infection, tumor cells were injected subcutaneously in the hind flank of nude mice. 6 days after injection, tumors size was measured daily for two weeks (Fig. 5F). We found that Mmg loss significantly slowed tumor growth starting from day 4 of measurement (Fig. 5G). At the end of the experiment, we evaluated Gli1 levels in randomly sampled tumors and found that Mmg loss significantly reduced Hh signal activity (Fig. 5H). Thus, knockdown of Mmg suppressed the growth of Hh-related tumors.

## Discussion

Genetic removal of PKA leads to full activation of the Hh pathway in the developing neural tube(Epstein *et al.*, 1996; Tuson, He and Anderson, 2011), suggesting PKA as a strong inhibitor of the Hh signaling. However, as a multifaceted enzyme, PKA is widely involved in many signaling and metabolic pathways. Therefore, global inhibition of PKA is not a feasible strategy for treatment. Our previous study pointed PDE4D as a potential target to inhibit Hh signaling(Ge *et al.*, 2015). Our current study highlights an effective approach to selectively inhibit PDE4D at one specific subcellular site. We provide evidence that dislocating PDE4D3 from the centrosome overactivates PKA locally at the centrosome to inhibits the Hh pathway, while sparing other PKA-related cellular events.

Cells have evolved two mechanisms to accurately govern local levels of cAMP and PKA activity: 1) controlling its production by adenylyl cyclase, and 2) managing its degradation by cAMP-specific PDEs. We believe that activation of Hh signaling involves both mechanisms. The first mechanism has been shown to be mediates by GPR161 that resides at the cilium. When the Hh pathway is off, GPR161 activates the Gαs-adenylyl cyclase pathway and keeps the local cAMP levels high(Mukhopadhyay *et al.*, 2013). Upon Shh stimulation, GPR161 exits the cilium and stops cAMP production(Mukhopadhyay *et al.*, 2013). Synergistic to this mechanism, PDE4D at the centrosome degrades the cAMP that is diffused from the nearby subcellular areas. The combined effects of these two mechanisms keep local PKA activities in check to allow the ensuing Hh signaling events to occur. When PDE4D activity is absent from the centrosome, the local cAMP concentration fails to reduce to the subthreshold level, even though the GPR161-Gαs-adenylyl cyclase pathway stops to produce cAMP. As a result, the high PKA levels at the centrosome suppresses the Hh signal transduction. Our study, for the first time, unmasked the roles of centrosomal PDE4D in the Hh pathway.

PDE4D is a large protein family comprising more than 12 alternative splicing isoforms in mammalian cells(Maurice *et al.*, 2014). PDE4D3 was originally identified to bind to Mmg by Verde et al (2001). Besides PDE4D3, other isoforms could be tethered to the centrosome as well, and Mmg loss might dislocate all these isoforms from the centrosome. It is also noteworthy that after adding IBMX, FRET was not completely abolished in Mmg knockout cells, indicating the existence of other PDE isoforms at the centrosome (Fig. 3G). It will be intriguing for future studies to delineate the identify of these PDE isoforms and their targeting mechanism to the centrosome.

Our study pointed to an effective method to suppress Hh signaling in cancers. Current Hh inhibitors target Smo, and these inhibitors are facing challenges of drug resistance and tumor relapse(Yauch *et al.*, 2009). Since the Mmg-PDE4D-PKA axis acts directly at Gli transcription factors, downstream of Smo, targeting this axis will be effective for cancers that have developed resistance to Smo inhibitors. Further, it is known that the basal activity of PDE4D is high, and most PDE4D small molecular inhibitors act by blocking the catalytic domain of PDE4D(Gavaldà and Roberts, 2013). These inhibitors block all PDE4D isoforms and are associated with severe side effects. Our results suggest that we may eliminate PDE4D activity specifically from the centrosome without blocking its catalytic domain. It pinpointed an effective therapeutic avenue to treat Hh-related cancers with reduced side effects.

## Materials and Methods

### Plasmids and generation of Myomegalin knockout CRISPR cell clones

Human PDE4D3 is generously provided by Marco Conti lab at UCSF, and was subcloned to include HA and EGFP tag. Myomegalin-C is cloned by RT-PCR with a mouse total mRNA library, and subcloned to included Flag and EGFP tag. The FRET probe AKAR4 was generously provided by Jin Zhang lab (available in Addgene). pcDNA3-mPKA-RIIα-AKAR4-NES was constructed by linking mPKA-RIIα with AKAR4-NES. The linker sequence is GGGGSGS. The two gRNA were designed via the Guide Design Resources of Feng Zhang lab at MIT(https://zlab.bio/guide-design-resources). The two gRNAs were cloned into the backbone of pX330-U6-Chimeric_BB-CBh-hSpCas9 (Addgene 42230), and transfected into MEF cells together with EGFP via lipofectamine 2000. 48hr after transfection, EGFP-positive cells were sorted by flow cytometer and plated into individual wells in 96-well plate. Individual cell clones were cultured for 2-4 weeks, and transferred to 24-well plate for further expansion.

### Time-lapse image with AKAR4

MEF cells were co-transfected with RIIα-AKAR4 and RFP-PACT via electroportation, and cultured in DMEM supplemented with 10% FBS at 37°C. 24hr later, cells were plated onto 8-chambered lab-Tex II coverglass (Thermo Fisher) at a density of 3.5 × 10^4^/well, and then grown for approximately 24h before imaging.

For imaging, cells were washed once with Extracellular Imaging Buffer (ECB, 5mM KCl, 125mM NaCl, 1.5mM CaCl_2_, 1.5mM MgCl_2_, 10mM Glucose, 20mM HEPES) and kept in ECB in the dark at room temperature. Images were collected with an epiflourescence microscope (Zeiss Observer3) with a 40X dry 0.9 NA objective lens connected to two linked Hamammatsu Flash v3 sCMOS cameras to facilitate real-time FRET imaging. The CFP fluorophore was excited using a 430 nm LED (Colibi7, Zeiss), and emission collected using a triple-bandpass emission filter, 467/24 + 555/25 + 687/145 (set 91 HE from Zeiss). Downstream, the collected emission was further split onto the two cameras using a 520 nm dichroic. Exposure time was set for 200ms. Images were acquired every 2min. IBMX was added to the cell as indicated in the experiment.

### FRET analysis

Results were analyzed in ImageJ. The centrosome was identified in red channel via RFP-PACT and selected as region of interest (ROI). An automated, stack-based thresholding was built on Renyi entropy method to identify strong fluorescence in the RFP channel throughout the time course. Intensities of the CFP and YFP at each time point in the ROI were measured. To control for different expression levels of AKAR, intensity at each time point was normalized to time zero.

### Western blot

Cells were lysed on ice in RIPA buffer containing 25mM Tris-HCl (pH7.6), 150mM Nacl, 1% NP-40, 1% sodium deoxycholate, 0.1% SDS, 1mM PMSF, 10mM sodium fluoride, 2mM sodium pyrophosphate, 1mM sodium orthovanadate, Roche protease inhibitor cocktail and Roche PhosSTOP inhibitor cocktail for 30 min. Lysates were cleared with centrifugation at 13,000 rpm for 30 min at 4℃. Protein concentrations of the supernatants were determined with BCA protein assay kit (Pierce). Protein samples were boiled in 6x SDS sample buffer for 10 min, and resolved in SDS-PAGE. Protein binds were transferred to PVDF membrane (88520, Thermofisher), which were blocked in Tris buffer (PH7.0) containing 0.1% Tween-20 and 5% BSA. The membrane was incubated in primary antibodies (diluted in blocking buffer) overnight at 4 ℃, and washed 3 times before incubation with HRP-conjugated secondary antibodies. Protein bands were visualized with ECL Western Blot substrate (Pierce, 32109).

Primary antibodies used: mouse anti-GAPDH (ab9484, Abcam), rabbit anti-phosphoPKA-T197 (5661S, Cell Siganling), rabbit anti-phosphoCREB-S133 (9198S, Cell Signaling), mouse anti-PKA (610625, BD Biosciences), rabbit anti-Gli1 (V812, Cell Signaling), rabbit anti-Myomegalin (PA5-30324, Invitrogen).

### Co-Immunoprecipitation

Plasmids were transfection into HEK293Tcells with lipofectamine 2000 reagent (Invitrogen) according to manufacturer’s instruction. Plasmids used : HA-PDE4D3, 3xFlag-Mmg-C560 and 3xFlag-vector (E4026, Sigma, MO). 24hr after transfection, cells were lysed in ELB buffer (150mM NaCl, 1% TritonX100, 50mM Tris pH8.0, 5mM EDTA, 5mM NaF, 2mM Na_3_VO_4_) supplemented with Protease inhibitor cocktail (Roche 11836170001) for 30min at 4°C. Lysates were cleared by centrifugation at 14,000rpm for 15min. Protein concentration of supernatants were determined using Pierce BCA Protein Assay Kit (Thermo Scientific). Equal amount of protein was loaded to the anti-Flag M2 Magnetic beads (M8823, Sigma) and anti-HA Magnetic beads (88836, Thermo Scientific) and incubated for 1h at room temperature. Beads were washed according to manufacturer’s instruction and incubated with 2x Laemmli sample buffer at 95°C for 5min. Samples were loaded to 10% SDS-PAGE gel and western blot was performed. Antibodies used: HA-tag rabbit antibody (3724S, Cell Signaling), anti-Flag M2 antibody (F1804, Sigma).

### Immunofluorescence staining

NIH3T3 or MEF cells grown on Poly-D-Lysine (A003E, Sigma) coated coverslips were fixed with 4% paraformaldehyde for 10min at room temperature. Cells were then blocked with 2% donkey serum and 0.1% triton in PBS for 1h. Primary and secondary antibodies are incubated with cells in the blocking buffer. Images were taken with LEICA DMi8 microscopy, or Zeiss LSM880 confocal microscope, with 60x oil lens.

Primary antibodies used: anti-Flag M2 antibody (F1804, Sigma), anti-mouse Pericentrin (611814, BD Biosciences), anti-rabbit myomegalin antibody (PA552969, Invitrogen), anti-rabbit GFP antibody (A11122, Thermo Fisher), mouse anti-acetylated tubulin (T6793, SIGMA), goat anti-Gli2 (AF3635, R&D SYSTEMS), rabbit anti-pPKA (ab59218, Abcam).

### Quantification of PhosphoPKA, PDE4D3 and Gli2

The levels of phosphoPKA and EGFP-PDE4D3 in centrosome were measured using ImageJ software as follows. First, an area of interest (AOI) was delineated based on the signal intensity of pericentrin staining; second, the mean gray value in AOI was measured in the phosphoPKA or EGFP-PD4D3 channel (F1); third, the contour of AOI was manually dragged to a nearby region within the cell, and the mean gray value of the enclosed area was measured as background (F2). The final values of phosphoPKA and PDE4D3 were calculated as F = F1-F2.

To quantify Gli2 levels at the cilium tips, the contour of the cilium tips was outlines in red channel (acetylated tubulin staining). The mean gray density in the enclosed area was measure in green channel (Gli2 staining). The background gray density was measured and subtracted to obtain the final Gli2 intensity at the cilium tips.

For each condition, 35-60 cells were measured. In myomegalin rescue experiment, only cells that were transfected with EGFP-Mmg are measured. Data analysis were done with Graphpad Prism 8.0 Software. Kruskal–Wallis non-parametric One-Way ANOVA was used for statistical analysis.

### Quantitative PCR

Cells were plated in 6-well plates at 0.5 × 10^6^ cells per well and cultured overnight. For Hh induction, cells were stimulated with 100nM SAG in starvation medium (0.5% FBS in DMEM) for 20-24hr. Total RNAs were isolated with Trizol reagent. The concentration of total RNA was normalized, and the same amount of RNA was mixed with qScript XLT-1 Step, RT-qPCR ToughMix (Quantabio 66149433,) together with specific TaqMan expression assays. The real-time PCR is performed in QuantStudio 3 (ThermoFisher).

The following TaqMan gene expression probes used: Mm00494654_g1 (Gli1), Mm00626240_m1 and Mm01257004_m1 (Mmg), Mm99999915_g1 (GAPDH).

### Primary culture of GNPs and Brdu incorporation assay

Neonate CD1 mouse were sacrificed at P7. The cerebellum was taken out and cut into small pieces with razor blades, incubated at 37°C for 15min in digestion buffer (HBSS with 20mM HEPES, PH 7.3, supplemented with trypsin and DNase I). At the end of incubation, digestion buffer was aspirated and replaced with Neurobasal medium with 250U/ml DNase I. Tissues were then triturated with pipet tips and polished Pasteur pipettes. After seated for 2min, dissociated cells were collected from the upper layer and centrifuged at 1000rpm for 5 min. Cells were washed one time, and resuspended in Neurobasal, supplemented with B27 (17504044, Gibco), Glutamax, and 1% Penicillin/streptomycin. Cells were then plated on coverslips coated with Poly-D-Lysine and Laminin. Lentivirus expressing Mmg shRNA were added 5hr after plating, and incubated overnight. 48hr after plating, 10nM SAG was added to the cells and incubated overnight. Brdu (20μM) was added to the culture and pulse labeled for 4hr, after which cells are fixed with 4% paraformaldehyde. Brdu immunostaining were performed with mouse-anti-Brdu antibody (662411Ig, Proteintech).

### Medulloblastoma xenograft mouse model

All the *in vivo* surgery steps and treatments were performed in accordance with the animal protocols approved by UC Merced’s Institutional Biosafety Committee (IACUC). MB56 tumor cells were cultured in neurobasal medium supplemented with B-27 (21103-049, Thermofisher). Cells were infected with lentiviral particles of shRNA against Myomegalin or control shRNA.

48hr later, cells were then collected, centrifuged and resuspended in PBS at 2 × 10^7^cells per 50ul. 50ul cells were mixed with Matrigel (354234, ThermoFisher) at 1:1 volume ration. The 100ul mixture was slowly injected into hind flanks of 8-10 nude mice of 7-week old (002019, Jackson Laboratory) under isoflurane anesthesia. 6-days after injection, the tumor volume was measured daily with digital caliper for two weeks. At the end of the experiments, mice were euthanized and 5-6 tumors were harvested randomly. Tumor tissues were proceeded to RNA extraction and qPCR. To generate a tumor growth curve, the relative tumor size is calculated as the ration of tumor size on each day over the size of the same tumor on day 1 of measurement.

### Quantification and Statistical Analysis

Statistical analysis was performed using Graphpad Prism 8.0 Software (Graphpad Software; La Jolla, CA, USA). Statistical significance was determined by Student’s t-Test or Kruskal–Wallis non-parametric One-Way ANOVA as mentioned in the figure legends.

## Acknowledgements

We thank Dr. Marco Conti for their generous gifts of PDE4D constructs, and helpful discussions. We thank Lavpreet Jammu, Anh Diep and Christi Waer for initial characterization of Myomegalin shRNA and for generating Myomegalin and PDE4D3 constructs. We thank Dr. Lin Gan for helpful discussions on the manuscript. The research was supported by N.I.H. grant R15 CA235749 to X.G.

## Author contributions

Jingyi Zhang: Acquisition, analysis and interpretation of data on Mmg-PDE4D interaction, GNP culture, and FRET assay

Hualing Peng: Acquisition, analysis and interpretation of data on characterization of Mmg CRISPR Knockout clones and mouse tumor model

Winston Ma: Establishment, expanding, initial selection and maintenance of Mmg CRISPR Knockout clones

Amanda Ya: Initial selection and maintenance of Mmg CRISPR Knockout clones, Characterization of INDEL mutations in Mmg CRISPR clones

Sammy Villa: Cloning and transfection of Mmg CRISPR construct into MEF cells

Shahar Sukenik: Supervision of the AKAR4 experiment and FRET data analysis

Xuecai Ge: Conception and design of all experiments, Acquisition of a portion of data, Analysis and interpretation of data, Drafting the article

---

**S1 (related to Figure 2).**
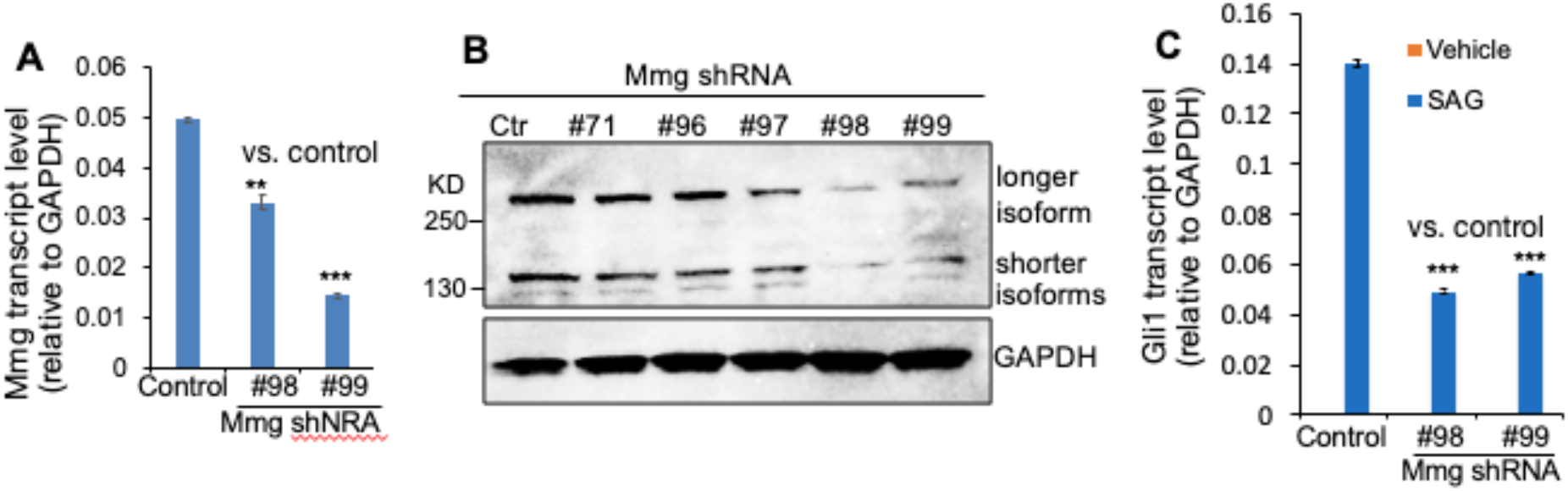
Mmg knockdown impairs Hh signal transduction. (A) NIH3T3 cells were transfected with shRNA against myomegalin. 72hr after transfection, myomegalin transcript levels were assessed by qPCR. Two of the shRNAs significantly reduced Mmg transcription. (B) Western blot showing that among all the shRNAs tested, #98 and #99 decreased the Mmg protein expression levels. (C) 72hr after shRNA transfection, NIH3T3 cells were treated with SAG overnight. Hh signaling activity was evaluated by qPCR to detect the transcript levels of Gli1, a Hh pathway target gene. All error bars represent SD; statistics: Student’s t-Test. **p<0.01, ***p<0.001.

**S2 (related to Fig 2).**
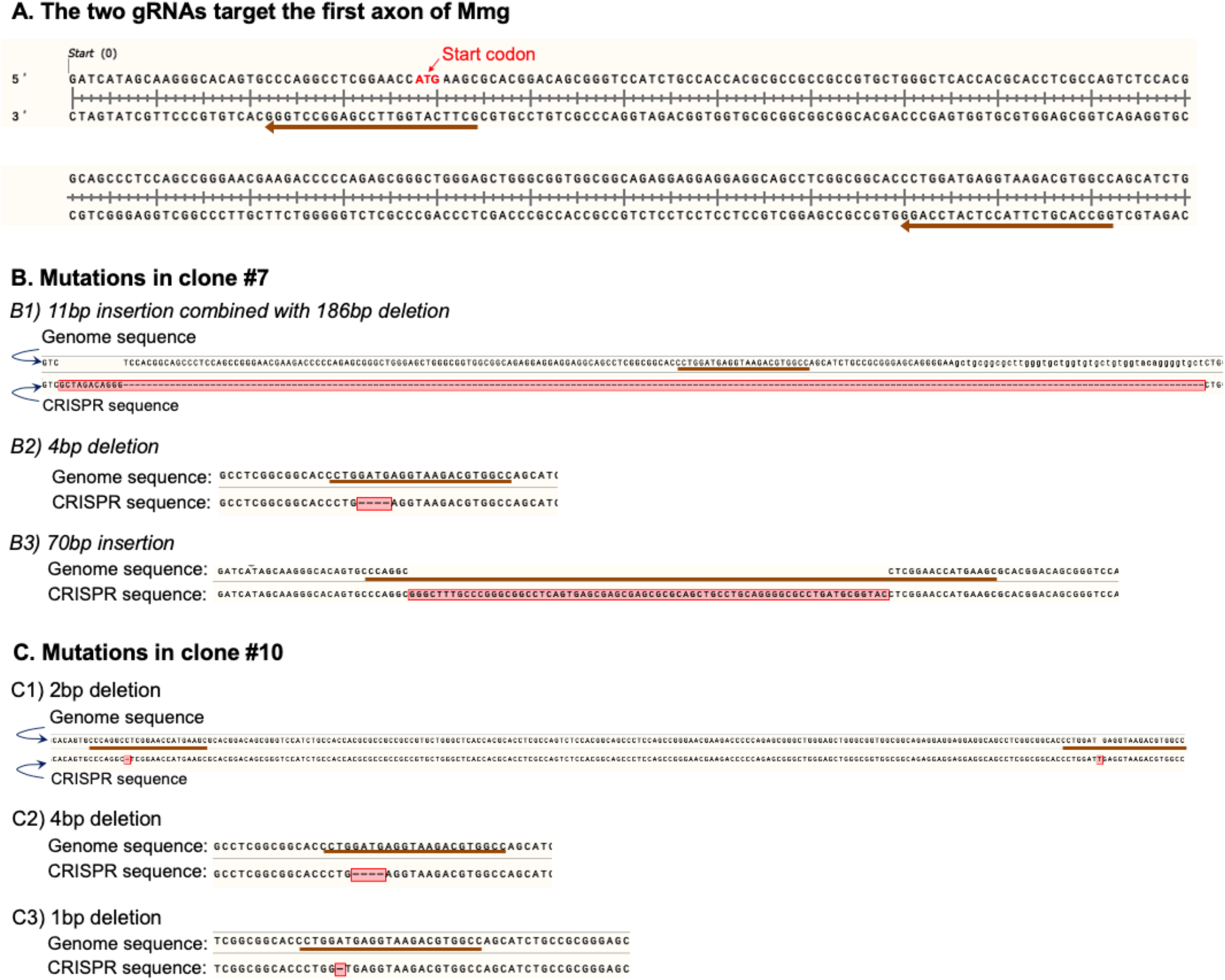
Design of Mmg CRISPR and genomic mutations in individual cell clones. (A) Design of Mmg CRISPR. Two guide RNAs were designed and both target exon 1 of mouse Mmg. Brown arrows underlie the sequence of gRNAs. (B-C) The gRNA targeting region in mouse genomic was amplified by genomic PCR, ligated into TOPO vector, and transfected into chemically competent cells. 20 bacterial colonies of each cell clones were randomly picked and sequenced. 3 type of mutations were found in cell line #7 (B) and #10 (C). No wild type sequences were identified in the 20 colonies, suggesting that all alleles of Mmg were mutated. Mutations lead to frame shift (B1, B2, C1, C2, C3), or alter the 5’ UTR (B3) that prevents the initiation of translation, all eventually leading to nonsense-mediated decay (NMD) of the mutant mRNA.

**S3 (related to Fig 3).**
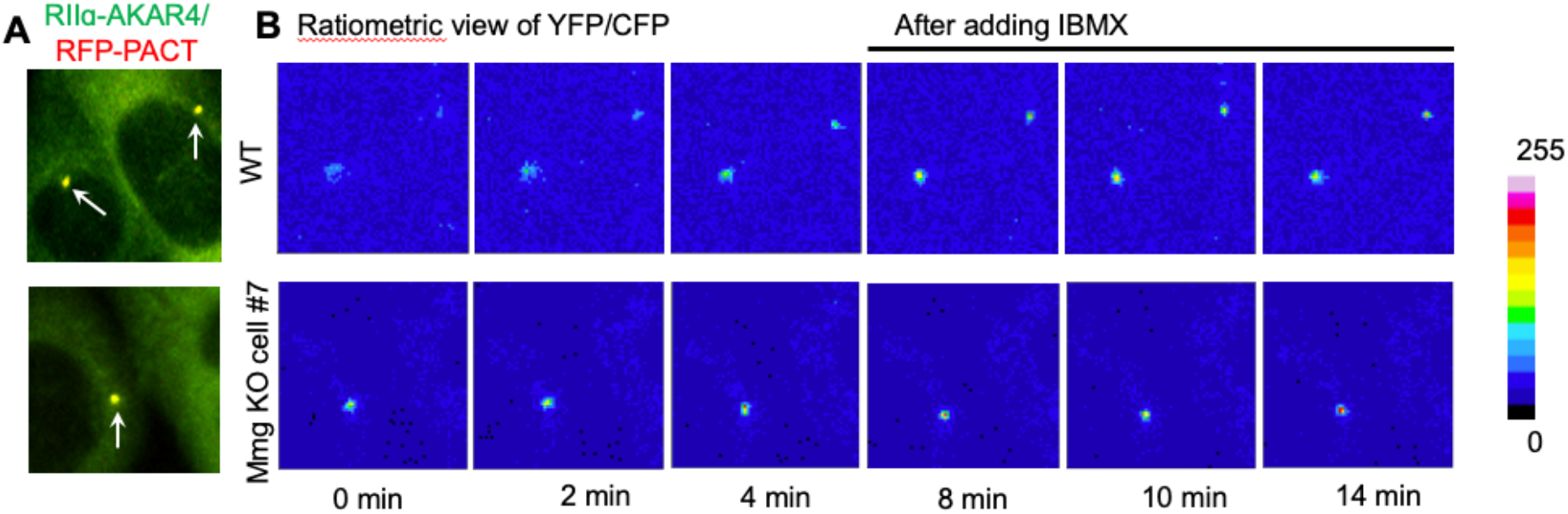
Mmg loss blocked IBMX’s effect on PKA activity at the centrosome. (A) Fluorescence images in live cells con-expressing RFP-PACT and RIIɑ-AKAP4. The centrosomes (white arrow) was identified by RFP-PACT in the red channel, selected as the region of interest (ROI), and FRET signaling were analyzed in the ROI. (B) Ratiometric view of FRET efficiency before and after IBMX treatment.

## References

Barzi, M. et al. (2010) ‘Sonic-hedgehog-mediated proliferation requires the localization of PKA to the cilium base’, Journal of Cell Science, 123(1), pp. 62–69. doi: 10.1242/jcs.060020.

Briscoe, J. and Therond, P. P. (2013) ‘The mechanisms of Hedgehog signalling and its rolesin developement and disease’, Nature Reviews Molecuar Cell Biology, 14(7), p. 416. doi: http://dx.doi.org.ezproxy.lib.ucalgary.ca/10.1038/nrm3598.

Dahmane, N. and Ruiz i Altaba, a (1999) ‘Sonic hedgehog regulates the growth and patterning of the cerebellum.’, Development (Cambridge, England), 126(14), pp. 3089–3100.

Epstein, D. J. et al. (1996) ‘Antagonizing cAMP-dependent protein kinase A in the dorsal CNS activates a conserved Sonic hedgehog signaling pathway.’, Development (Cambridge, England), 122, pp. 2885–2894.

Fouladi, M. et al. (2005) ‘Intellectual and functional outcome of children 3 years old or younger who have CNS malignancies’, Journal of Clinical Oncology, pp. 7152–7160. doi: 10.1200/JCO.2005.01.214.

Gavaldà, A. and Roberts, R. S. (2013) ‘Phosphodiesterase-4 inhibitors: a review of current developments (2010 - 2012).’, Expert opinion on therapeutic patents, 23(8), pp. 997–1016. doi: 10.1517/13543776.2013.794789.

Ge, X. et al. (2015) ‘Phosphodiesterase 4D acts downstream of Neuropilin to control Hedgehog signal transduction and the growth of medulloblastoma’, eLife. eLife Sciences Publications Ltd, 4(September2015).

Gillingham, A. K. and Munro, S. (2000) ‘The PACT domain, a conserved centrosomal targeting motif in the coiled-coil proteins AKAP450 and pericentrin’, EMBO Reports, 1(6), pp. 524–529. doi: 10.1093/embo-reports/kvd105.

Goodrich, L. V. et al. (1997) ‘Altered neural cell fates and medulloblastoma in mouse patched mutants’, Science. doi: 10.1126/science.277.5329.1109.

Han, Y.-G. and Alvarez-Buylla, A. (2010) ‘Role of Primary Cilia in Brain Development and Cancer.’, Current opinion in Neuroviology, 20(1), pp. 58–67. doi: 10.1016/j.conb.2009.12.002.Role.

Herbst, K. J., Allen, M. D. and Zhang, J. (2011) ‘Spatiotemporally Regulated Protein Kinase A Activity Is a Critical Regulator of Growth Factor-Stimulated Extracellular Signal-Regulated Kinase Signaling in PC12 Cells’, Molecular and Cellular Biology, 31(19), pp. 4063–4075. doi: 10.1128/mcb.05459-11.

Houslay, M. D. (2010) ‘Underpinning compartmentalised cAMP signalling through targeted cAMP breakdown’, Trends in Biochemical Sciences, pp. 91–100. doi: 10.1016/j.tibs.2009.09.007.

Huang, Y., Roelink, H. and McKnight, G. S. (2002) ‘Protein kinase A deficiency causes axially localized neural tube defects in mice’, Journal of Biological Chemistry, 277(22), pp. 19889–19896. doi: 10.1074/jbc.M111412200.

Hui, C. and Angers, S. (2011) ‘Gli Proteins in Development and Disease’, Annual Review of Cell and Developmental Biology, 27(1), pp. 513–537. doi: 10.1146/annurev-cellbio-092910-154048.

Humke, E. W. et al. (2010) ‘The output of Hedgehog signaling is controlled by the dynamic association between Suppressor of Fused and the Gli proteins’, Genes and Development. doi: 10.1101/gad.1902910.

Kool, M. et al. (2012) ‘Molecular subgroups of medulloblastoma: An international meta-analysis of transcriptome, genetic aberrations, and clinical data of WNT, SHH, Group 3, and Group 4 medulloblastomas’, Acta Neuropathologica, 123(4), pp. 473–484. doi: 10.1007/s00401-012-0958-8.

Maurice, D. H. et al. (2014) ‘Advances in targeting cyclic nucleotide phosphodiesterases.’, Nature reviews. Drug discovery. Nature Publishing Group, 13(4), pp. 290–314. doi: 10.1038/nrd4228.

McCormick, K. and Baillie, G. S. (2014) ‘Compartmentalisation of second messenger signalling pathways’, Current Opinion in Genetics and Development. Elsevier Ltd, 27, pp. 20–25. doi: 10.1016/j.gde.2014.02.001.

Mukhopadhyay, S. et al. (2013) ‘The ciliary G-protein-coupled receptor Gpr161 negatively regulates the sonic hedgehog pathway via cAMP signaling’, Cell, 152(1-2), pp. 210–223. doi: 10.1016/j.cell.2012.12.026.

Purzner, T. et al. (2018) ‘Developmental phosphoproteomics identifies the kinase CK2 as a driver of Hedgehog signaling and a therapeutic target in medulloblastoma’, Science Signaling. doi: 10.1126/scisignal.aau5147.

Richter, W., Jin, S.-L. C. and Conti, M. (2005) ‘Splice variants of the cyclic nucleotide phosphodiesterase PDE4D are differentially expressed and regulated in rat tissue.’, The Biochemical journal, 388(Pt 3), pp. 803–811. doi: 10.1042/BJ20050030.

Roubin, R. et al. (2013) ‘Myomegalin is necessary for the formation of centrosomal and Golgi-derived microtubules’, Biol Open, 2(2), pp. 238–250. doi: 10.1242/bio.20123392.

Shaywitz, A. J. and Greenberg, M. E. (1999) ‘CREB: A Stimulus-Induced Transcription Factor Activated by A Diverse Array of Extracellular Signals’, Annual Review of Biochemistry. doi: 10.1146/annurev.biochem.68.1.821.

Somatilaka, B. N. et al. (2020) ‘Ankmy2 Prevents Smoothened-Independent Hyperactivation of the Hedgehog Pathway via Cilia-Regulated Adenylyl Cyclase Signaling’, Developmental Cell. Elsevier Inc., 54(6), pp. 710–726.e8. doi: 10.1016/j.devcel.2020.06.034.

Tukachinsky, H., Lopez, L. V. and Salic, A. (2010) ‘A mechanism for vertebrate Hedgehog signaling: Recruitment to cilia and dissociation of SuFu-Gli protein complexes’, Journal of Cell Biology. doi: 10.1083/jcb.201004108.

Tuson, M., He, M. and Anderson, K. V. (2011) ‘Protein kinase A acts at the basal body of the primary cilium to prevent Gli2 activation and ventralization of the mouse neural tube’, Development, 138(22), pp. 4921–4930. doi: 10.1242/dev.070805.

Tyler Hillman, R. et al. (2011) ‘Neuropilins are positive regulators of Hedgehog signal transduction’, Genes and Development. doi: 10.1101/gad.173054.111.

Verde, I. et al. (2001) ‘Myomegalin is a Novel Protein of the Golgi/Centrosome That Interacts with a Cyclic Nucleotide Phosphodiesterase’, Journal of Biological Chemistry, 276(14), pp. 11189–11198. doi: 10.1074/jbc.M006546200.

Wallace, V. A. (1999) ‘Purkinje-cell-derived Sonic hedgehog regulates granule neuron precursor cell proliferation in the developing mouse cerebellum’, Current Biology, 9(8), pp. 445–448. doi: 10.1016/S0960-9822(99)80195-X.

Wang, B. and Li, Y. (2006) ‘Evidence for the direct involvement of βTrCP in Gli3 protein processing’, Proceedings of the National Academy of Sciences of the United States of America, 103(1), pp. 33–38. doi: 10.1073/pnas.0509927103.

Wechsler-Reya, R. J. and Scott, M. P. (1999) ‘Control of neuronal precursor proliferation in the cerebellum by sonic hedgehog’, Neuron, 22(1), pp. 103–114. doi: 10.1016/S0896-6273(00)80682-0.

Williams, C. H. et al. (2015) ‘An In Vivo Chemical Genetic Screen Identifies Phosphodiesterase 4 as a Pharmacological Target for Hedgehog Signaling Inhibition’, Cell Reports, 11(1), pp. 43–50. doi: 10.1016/j.celrep.2015.03.001.

Yamanaka, H. et al. (2010) ‘Forskolin, a Hedgehog signal inhibitor, inhibits cell proliferation and induces apoptosis in pediatric tumor cell lines’, Molecular Medicine Reports. doi: 10.3892/mmr-00000230.

Yamanaka, H. et al. (2011) ‘Hedgehog signal inhibitor forskolin suppresses cell proliferation and tumor growth of human rhabdomyosarcoma xenograft’, Journal of Pediatric Surgery. doi: 10.1016/j.jpedsurg.2010.11.010.

Yauch, R. L. et al. (2009) ‘Smoothened mutation confers resistance to a Hedgehog pathway inhibitor in medulloblastoma.’, Science (New York, N.Y.), 326(5952), pp. 572–4. doi: 10.1126/science.1179386.

Zaccolo, M. and Pozzan, T. (2002) ‘Discrete microdomains with high concentration of cAMP in stimulated rat neonatal cardiac myocytes’, Science, 295(5560), pp. 1711–1715. doi: 10.1126/science.1069982.

Zhang, J. et al. (2001) ‘Genetically encoded reporters of protein kinase A activity reveal impact of substrate tethering’, Proceedings of the National Academy of Sciences, 98(26), pp. 14997–15002. doi: 10.1073/pnas.211566798.

